# Individual boldness is life stage-dependent and linked to dispersal in a hermaphrodite land snail

**DOI:** 10.1101/098822

**Authors:** Maxime Dahirel, Alexandre Vong, Armelle Ansart, Luc Madec

## Abstract

Both individual variation in dispersal tendency and animal personalities have been shown to be widespread in nature. They are often associated in personality-dependent dispersal, and both have major but underappreciated consequences for ecological and evolutionary dynamics. In addition, personalities are not stable over time and changes can appear through ontogeny, leading to life stage-dependent behaviours. We investigated relationships between dispersal, life stage and boldness in an invertebrate with between- and within-life stages variation in dispersal tendency, the land snail *Cornu aspersum.* Latency to exit the shell following a simulated attack was repeatable, indicating boldness is a personality trait in *Cornu aspersum*. Subadults were bolder and more dispersive than adults. Dispersers were bolder than non-dispersers, independently of boldness changes between life stages. We discuss how these results can be explained in relation with life history strategies in this hermaphrodite species, in particular risk management in the context of reproductive investment.

## INTRODUCTION

Individual variation in dispersal, *i.e.* in movement leading to gene flow in space, is often correlated with variation in other phenotypic traits (life-history, physiology, morphology, behaviour…)[1–3]. Such “dispersal syndromes” may help dispersing and resident individuals to maximize fitness, offsetting costs incurred during dispersal and in the origin habitat, respectively [1]. Knowledge of these syndromes can yield insights on the proximate and ultimate mechanisms underlying dispersal decisions, and help better appreciate the consequences of dispersal on meta-population functioning [1].

Inter-individual differences in behaviour that are consistent across time or contexts (animal personalities) have been described for a large set of taxa and behaviours (although mostly in vertebrates; [4–8] and references therein). They have important yet under-assessed ecological and evolutionary consequences, as individuals exhibiting different personalities also often differ in various other life-history traits [6,8]. Boldness/shyness, broadly defined as an individual’s reaction to risky situations (e.g. predators) [4] is a key personality trait in the context of dispersal [3]. As dispersal presents many costs and risks, bolder individuals are often more dispersive [3]. Dispersal is also often age-structured, with the nature of the most dispersive life stage depending on age-dependent dispersal costs or trade-offs with reproductive investment [9]. While boldness changes through ontogeny have also been documented [10,11], we do not currently know to which extent dispersal-boldness syndromes are linked to age-/life stage-dependent dispersal.

Here we analyzed the relationships between boldness, dispersal and life stage in an invertebrate, the land snail *Cornu aspersum* (Müller) (family Helicidae). We tested whether boldness varied with life stage (subadult/ adult) and dispersal status in this species, to determine whether these differences could explain previously documented stage-dependent dispersal patterns [12,13].

## MATERIALS AND METHODS

### Snail collection and maintenance

In April 2016, snails were collected by hand in peri-urban parks in Rennes, France. Sixty adults and sixty subadults (greater shell diameter > 20 mm) were used in experiments; adults were recognisable by the presence of a lip around the shell peristome. Snails were individually marked with felt-tip paint markers and maintained under controlled conditions (20°C ± 1°C; 16L: 8D, with scotophase starting at 8:00 pm; *ad libitum* access to cereal flour-based snail food from Hélinove, Le Boupère, France). They were housed by groups of either 10 adults or 10 subadults in polyethylene boxes covered by a net (30 × 45 × 8 cm) and lined by synthetic foam kept saturated with water. Boxes were cleaned and the lining changed once per week. Two snails that died during the experiments were replaced by adding one new individual to two of the following boxes.

### Boldness tests

Our protocol is inspired by Seaman and Briffa [7]. Snails were assayed between 16:00 and 20:00, i.e. at the end of photophase. Snails were first placed in a Petri dish containing water for up to 5 minutes to stimulate activity, and then placed on glass plates. Once they moved at least 3 cm away from their starting position, an operator used a pipette tip to pinch them for 5 seconds on the right side of the foot, close to the peristome. This caused all snails to retract fully within their shells. All tests were carried out by the same operator (A.V.). We used the time snails took to exit their shells following the “attack”, from retraction to the full extension of all tentacles out of the shell, as our measure of boldness (snails with shorter latencies being considered bolder). Two trials, 7 days apart, were conducted on each snail. We tested all snails coming from the same box on the same day, and replaced them in their source box after testing. We stopped tests if a snail had not moved after 20 minutes; these interrupted observations (8 out of 240) were not included in further analyses.

### Dispersal tests

Dispersal was assessed in an outdoor tarmacked area with no food or shelter on the Beaulieu university campus, Rennes. Seven days after their second boldness test, snails were placed by life-stage in open boxes, around the middle of the arena for one night (19:00 to 09:00 on the following day). All tests were made on nights with mean temperature > 10 °C and rainfall ≤ 1 mm. We tested two boxes per night, one per life stage (adult/ subadult). Boxes tested the same night were separated by at least 6 m, a distance larger than this species’ perceptual range [13]. No dispersing snail was found in the other box the following morning. Both food and the box lining were left in boxes during tests in order to provide snails with a favourable and familiar environment, and one slate was added in each box to provide shelter. Based on available information on home range and routine movements, only snails recaptured farther than 1 m from the centre of their box were deemed dispersers [12].

### Statistical analyses

Analyses were done using R version 3.3.1 and the *lme4* package [14,15]. Differences in dispersal probability between subadults and adults were assessed using a binomial generalized linear mixed model, with a random effect of test night. Log-transformed latencies to resume activity were analysed using a linear mixed model with dispersal status, developmental stage and their interaction as explanatory factors, as well as random effects of individual identity and test session. We used the Satterthwaite approximation as implemented in the *ImerTest* package to determine degrees of freedom for conditional *F* tests [16]. We calculated mixed-model repeatabilities (raw and adjusted for the effects of model variables) and their 95% confidence intervals following Nakagawa and Schielzeth [17] using the *rptR* package. Repeatabilities significantly higher than zero indicate within-individual consistency in behaviour between trials, and were deemed evidence of animal personality [4].

## RESULTS

Both raw and adjusted repeatabilities were significantly greater than zero (*R_raw_* = 0.52, *SE* = 0. 07, 95% CI = [0.37, 0.64]; *R_adj_* = 0.48, *SE* = 0.07, 95% CI = [0.32, 0.60]).

Subadults were more likely to disperse (Wald test, *X^2^* = 4.094, *p* = 0.043, Fig. 1), and were significantly faster to exit their shells than adults (*F*_1,112.8_ = 11.038, *p* = 0.001; Fig. 2, left). Stage being equal, snails that dispersed were also bolder than those that did not (*F*_1,112.8_ = 3.970; *p* = 0.049; Fig. 2, right). There was no significant stage × dispersal status interaction (*F*_1,112.8_ = 0.331, *p* = 0.566).

**Figure 1.**
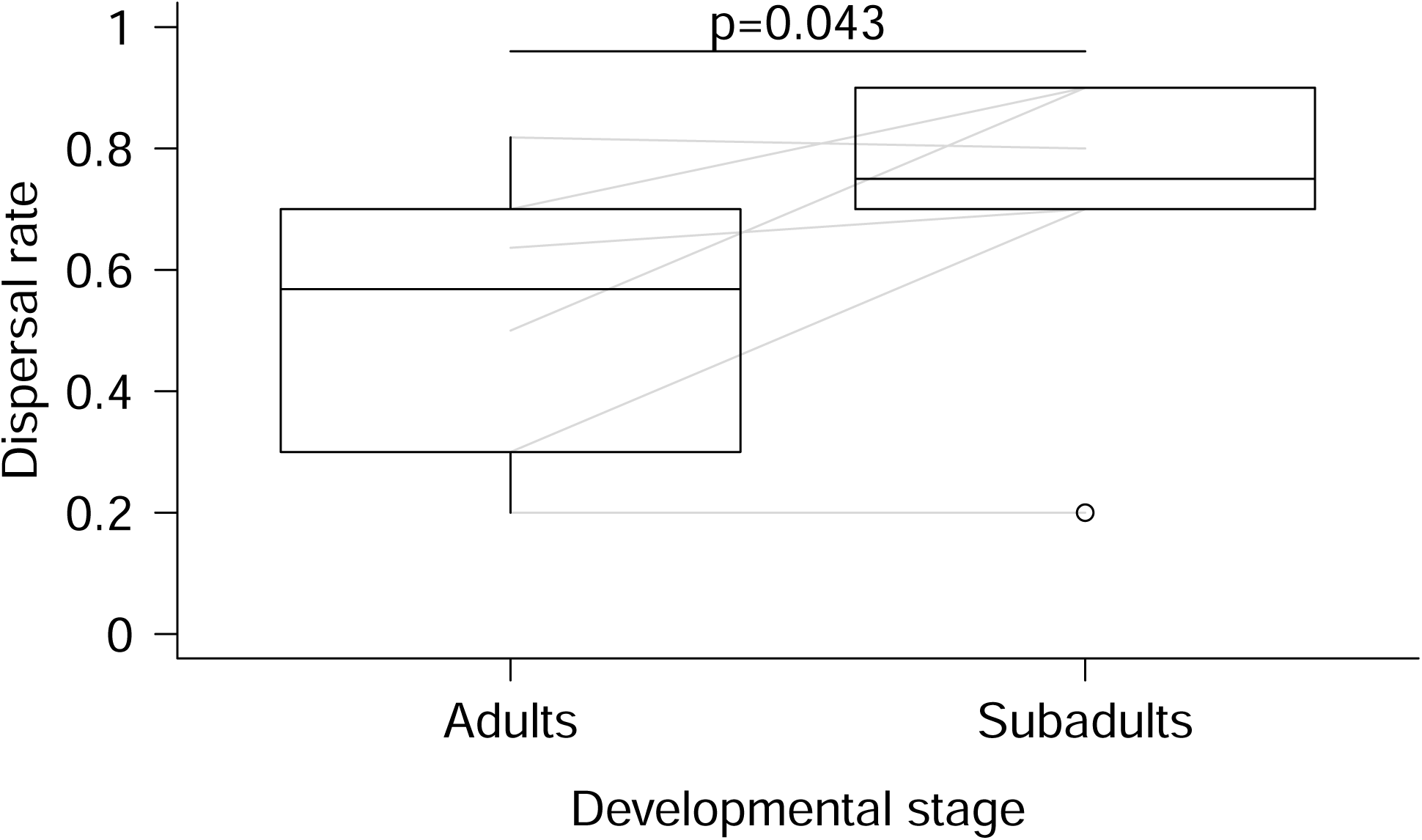
Dispersal rate per test box as a function of developmental stage. *P* value is based on a binomial generalized linear mixed model. Grey lines connect observed values of boxes tested on the same night.

**Figure 2.**
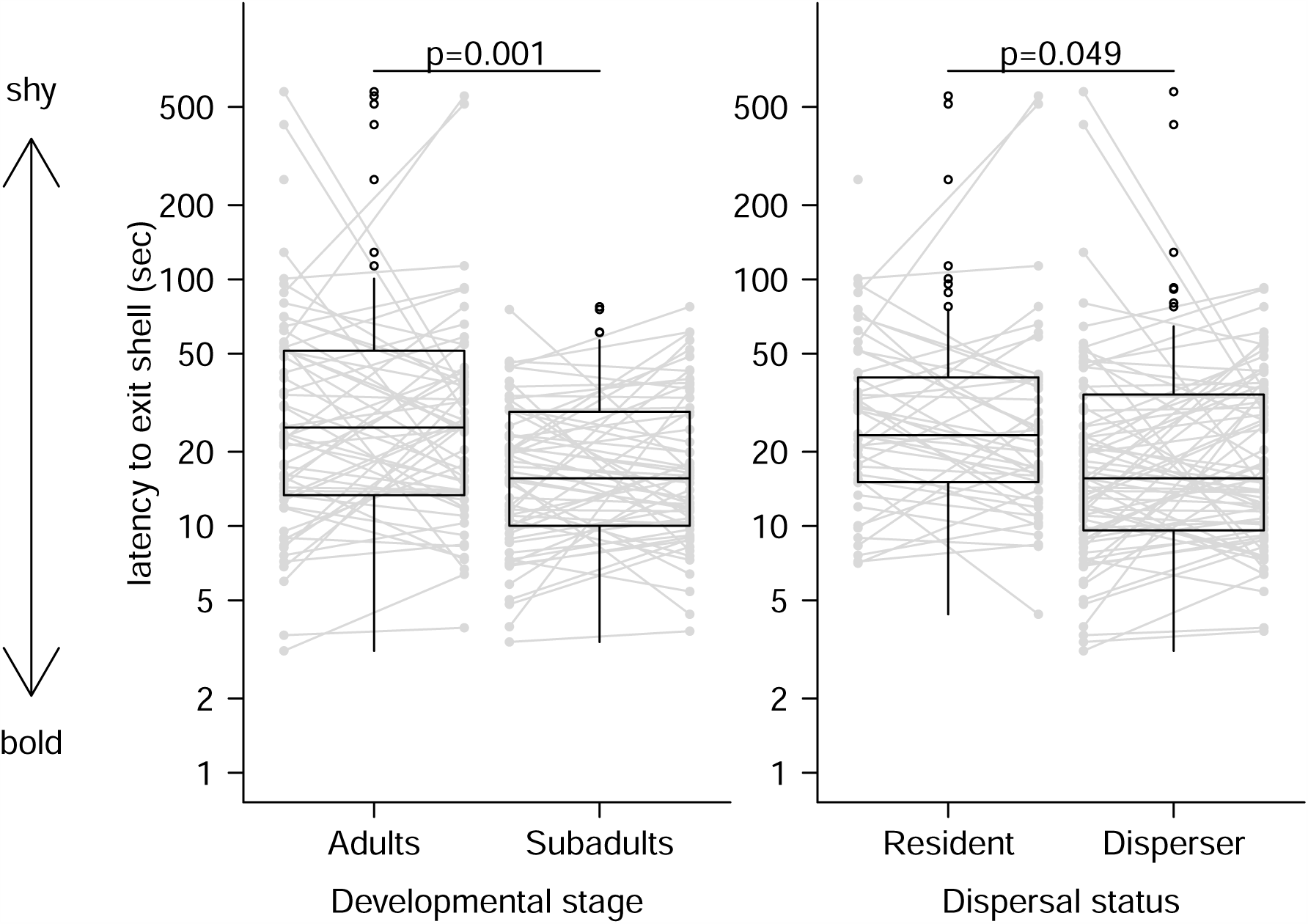
Stage- (left) and dispersal- (right) related differences in boldness (latency to exit shell). *P* values are based on a linear mixed model. Model was fitted, and data are plotted, on the log scale. Grey lines connect values from the same individual.

## DISCUSSION

Latency to exit the shell following a stressful stimulus was repeatable in *Cornu aspersum*, even after accounting for two potential confounding variables susceptible of increasing between-individual differences, namely life stage and dispersal status. The present study therefore provides, to our knowledge, the first evidence for animal personality in a terrestrial gastropod, following recent examples in aquatic species [7]. It adds to increasing evidence that animal personalities are widespread, even among non-vertebrates [5].

Subadults were more dispersive than adults (Fig. 1), a result in line with previous studies [12] and likely linked to the higher reproductive costs incurred by older snails [18]. Subadults were also bolder than adults (Fig. 2), and dispersers bolder than non-dispersers (Fig. 2). There was no interaction between life stage and dispersal status, meaning that in *Cornu aspersum*, personality-dependent dispersal is consistent across life stages, *i.e.* does not vary whether we consider the least (adult) or the most (subadult) dispersive life stage.

Bolder individuals are expected to have a lower survival on average, due to predation for instance [8]. Therefore, observed boldness differences between wild-caught subadults and adults might not reflect true behavioural shifts across ontogeny, but merely be the result of increased mortality of bold subadults, e.g. during dispersal [2,8]. While we were not able to separate these two effects here due to the use of wild individuals, true loss of boldness with aging/maturity has been observed in several other species [10,11]. Such behavioural shifts are expected when environmental situations and/ or life history expectations differ between life stages, with for instance later life stages being more risk-averse as a way to protect already acquired resources [11,19]. We expect these differences in life history expectations to play a major role in *Cornu aspersum*: although both stages are able to mate, adults are characterized by a large increase in female reproductive investment [20]. In addition, the same reasoning can be applied to explain the differences in boldness between dispersing and resident snails that were observed even after accounting for the effect of life stage. Indeed, dispersing *C. aspersum* snails present lower values of female investment than their more sedentary conspecifics [18]. This suggests a role of neuroendocrine factors associated with reproductive development [21] as common proximate drivers of both boldness and dispersal variation in this species [3].

*Cornu aspersum*’s recent history is characterized by serial introductions/ colonizations worldwide [22], and it is often present in highly anthropogenic fragmented environments [13]. Although questions remain on how the described boldness-dispersal syndrome changes with environmental context, and on the mechanisms underlying the life-stage dependency of boldness, we provide here some new information on the ecological consequences of individual personalities in an invertebrate species. Given the predicted role of personality-dependent dispersal in biological invasions and metapopulation dynamics [3], our results may help shed light on the mechanisms behind this species’ worldwide success.

## DATA ACCESSIBILITY

Data will be uploaded to Dryad or similar repositories upon acceptance.

## ACKNOWLEDGEMENTS

We thank Youn Henry for comments on a previous version of the manuscript.

## AUTHORS’ CONTRIBUTIONS

M.D., A.A. and L.M. conceived the study and designed the experiments. A.V. carried out the experiments. M.D. and A.V. conducted analyses and drafted the manuscript, with input from all authors. All authors gave final approval for publication.

